# A versatile new ubiquitin detection and purification tool derived from a bacterial deubiquitylase

**DOI:** 10.1101/2021.12.02.470885

**Authors:** Mengwen Zhang, Jason M. Berk, Adrian B. Mehrtash, Jean Kanyo, Mark Hochstrasser

## Abstract

Protein ubiquitylation is an important post-translational modification affecting an wide range of cellular processes. Due to the low abundance of ubiquitylated species in biological samples, considerable effort has been spent on developing methods to purify and detect ubiquitylated proteins. We have developed and characterized a novel tool for ubiquitin detection and purification based on OtUBD, a high-affinity ubiquitin-binding domain derived from an *Orientia tsutsugamushi* deubiquitylase. We demonstrate that OtUBD can be used to purify both monoubiquitylated and polyubiquitylated substrates from yeast and human tissue culture samples and compare their performance with existing methods. Importantly, we found conditions for either selective purification of covalently ubiquitylated proteins or co-isolation of both ubiquitylated proteins and their interacting proteins. As a proof-of-principle for these newly developed methods, we profiled the ubiquitylome and ubiquitin-associated proteome of the yeast *Saccharomyces cerevisiae*. Combining OtUBD affinity purification with quantitative proteomics, we identified potential substrates for E3 ligases Bre1 and Pib1. OtUBD provides a versatile, efficient, and economical tool for ubiquitin researchers with specific advantages over other methods, such as in detecting monoubiquitylation or ubiquitin linkages to noncanonical sites.

## Introduction

Ubiquitin is a highly conserved 76-residue protein present in all eukaryotic organisms (1). It is a post-translational protein modifier that requires a cascade of enzymatic reactions for its attachment to proteins. Each modification is catalyzed by a ubiquitin-activating enzyme E1, ubiquitin-conjugating enzyme E2, and ubiquitin ligase E3(2). The E1 enzyme activates the C-terminal carboxylate of ubiquitin by formation of E1-ubiquitin thioester. The E1 enzyme then transfers the activated ubiquitin molecule to an E2, with which it also forms a thioester linkage. E3 enzymes are responsible for the recognition of the substrate and catalyzing ubiquitin transfer from the E2 to a nucleophilic residue on the substrate protein, typically the ε-amino group of a lysine residue, but potentially also N-terminal amino groups, serine/threonine hydroxyl side chains, or the thiol group of cysteine (3). Ubiquitin itself can be ubiquitylated through its N-terminal methionine (M1) or one or more of its seven lysine residues (K6, K11, K27, K29, K33, K48, K63) (4). This enables diverse ubiquitin chain topologies and sizes, which modulate the biological functions of substrate ubiquitylation, often described as the “ubiquitin code” (5). For example, monoubiquitylation has been reported to facilitate protein complex formation in many cases (6, 7), polyubiquitylation involving K48 linkages is a well-documented substrate mark for proteasomal degradation (8), while polyubiquitylation with K63 linkages is often a signal for membrane trafficking or DNA repair pathways (9, 10). Ubiquitylation can be reversed through hydrolysis by a family of ubiquitin-specific proteases or deubiquitylases (DUBs) (11).

Defects in ubiquitylation have been connected to many human disorders, including cancers, viral infections and neurodegenerative diseases (12–14). The broad biomedical impact of protein ubiquitylation has stimulated efforts to develop sensitive methods to study the ubiquitylated proteome (15, 16). Because the ubiquitylated fraction of a given protein substrate population is often very small at steady state (17), it is generally necessary to enrich for the ubiquitylated proteins in biological samples of interest. Current methods to enrich ubiquitylated proteins can be roughly classified into three categories: 1) ectopic (over)expression of epitope-tagged ubiquitin and affinity purification using the tags, 2) immunoprecipitation with anti-ubiquitin antibodies and 3) use of tandem ubiquitin-binding entities (TUBEs) (18–23).

The first method was introduced using the yeast *Saccharomyces cerevisiae* (24). In yeast, four different genes encode ubiquitin, either as fusions to ribosomal peptides or as tandem ubiquitin repeats (25). It is possible to create yeast strains where the only source of ubiquitin is a plasmid expressing epitope-tagged ubiquitin (26, 27); as a result, all ubiquitylated proteins in this specific yeast strain bear the epitope tag, which can then be used for enrichment or detection of the ubiquitylated species. A number of earlier studies have used this method to profiled the ubiquitylated proteome (4). One major concern with this method is that the (over)expression of tagged ubiquitin may result in artifactual ubiquitylation or interfere with endogenous ubiquitylation events.

To study endogenous ubiquitylated proteins, anti-ubiquitin antibodies — including those against all ubiquitylation types (such as FK1 and FK2 monoclonal antibodies; (28)) or those specific for certain ubiquitin-chain linkages (such as anti-K48 ubiquitin linkage antibodies) — have been used (29, 30). TUBEs, on the other hand, are recombinant ubiquitin-affinity reagents built from multiple ubiquitin-binding domains (UBDs). UBDs have been characterized in a range of ubiquitin-interacting proteins, and they typically bind to ubiquitin with low affinity (31). By fusing multiple copies of a UBD together to turn it into a TUBE, the avidity of the reagent toward polyubiquitin chain-modified proteins is greatly increased (32). TUBEs are therefore useful in protecting polyubiquitylated proteins from DUB cleavages and enriching them in biological samples, and some TUBEs are designed to recognize specific types of polyubiquitin chains (33). In general, TUBE affinity toward monoubiquitylated proteins is low (32).

In addition to the above-mentioned methods, ubiquitin remnant motif antibodies (diGly antibodies) are widely used in bottom-up proteomics experiments to identify ubiquitylation sites on substrate proteins (34, 35). In bottom-up proteomics, proteins are digested by a protease (typically trypsin) into short peptides, separated by liquid chromatography and identified by tandem mass spectrometry (LC-MS/MS) (36). Tryptic digestion of ubiquitylated proteins leaves a signature GlyGly (GG) remnant on ubiquitylated lysine side chains (18). Anti-diGly-ε-Lys antibodies recognize this remnant motif and enrich such peptides for identification of ubiquitylation sites. The development of diGly antibodies has greatly facilitated the systematic discovery and profiling of ubiquitylated proteins and their ubiquitylation sites and has enabled the establishment of databases documenting ubiquitylation in humans and other species (37, 38).

Each of these methods has its advantages and limitations, which have been reviewed elsewhere (16, 39). For example, TUBEs are excellent tools to study polyubiquitylation, but in some mammalian cell types, over 50% of ubiquitylated proteins are only monoubiquitylated (17) and can easily be missed by TUBEs. Anti-diGly antibodies, while extremely effective in identifying ubiquitin-lysine linkages, are not capable of recognizing ubiquitylation sites on other nucleophilic side chains in proteins or other macromolecules (40). Due to the importance and complexity of ubiquitylation, the development of sensitive and economical reagents to study the entire ubiquitylome is crucial.

Recently, our group discovered a novel UBD within a deubiquitylase effector protein, OtDUB, from the intracellular bacterium *Orientia tsutsugamushi*, the causative agent of the disease scrub typhus (41). The UBD from OtDUB, which was referred to as OtDUB_UBD_ (we will use OtUBD for the remainder of the paper for simplicity), spans residues 170-264 of the 1369-residue OtDUB polypeptide (Fig. 1A) and binds monomeric ubiquitin with very high affinity (K_d_, ∼5 nM), which is more than 500-fold tighter than any other natural UBD described to date. Co-crystal structures of OtDUB and ubiquitin revealed that OtUBD binds ubiquitin at the isoleucine-44 hydrophobic patch, a ubiquitin feature commonly recognized by ubiquitin-binding proteins (42). We reasoned that the small, well-folded OtUBD could serve as a facile enrichment reagent for ubiquitylated proteins. The advantages of such reagent include its low cost, lack of bias between monoubiquitylated and polyubiquitylated proteins, and ability to detect unconventional ubiquitin-substrate linkages.

**Figure 1.**
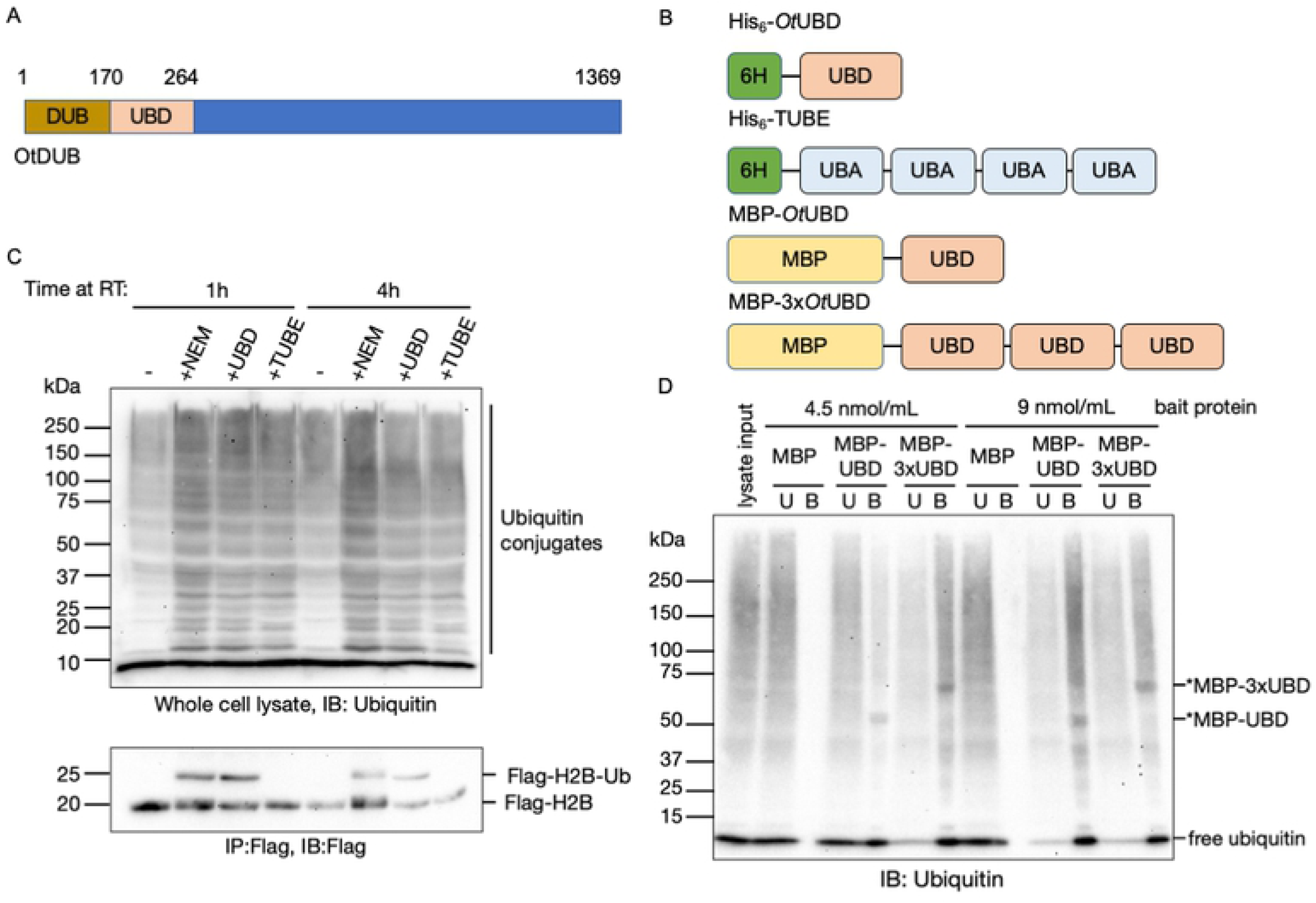
The high-affinity UBD from OtDUB efficiently protects and enriches for yeast ubiquitylated species. A. Schematic showing the ubiquitin binding domain (OtUBD) within the *O. tsutsugamushi* DUB (OtDUB). OtUBD spans residues 170 to 264.
B. The different constructs of OtUBD and the control TUBE derived from the UBA domain of human UBQLN1. His_6_ tagged OtUBD and TUBE were used in the ubiquitylation protection experiment shown in Fig. 1C and MBP-tagged OtUBD and 3xOtUBD were used in the ubiquitin pulldown experiment in Fig. 1D.
C. Immunoblot (IB) analysis of bulk proteins (top panel) and histone H2B (bottom panel) from yeast cell lysates prepared in the presence of different reagents. OtUBD prevents deubiquitylation of bulk ubiquitylated substrates (top panel) and monoubiquitylated histone H2B (bottom panel).
D. Immunoblot analysis of MBP pulldowns from yeast cell lysates using different bait proteins. MBP or MBP-tagged bait proteins bound to an amylose resin were incubated with yeast lysates, and bound proteins were eluted by incubation in SDS sample buffer. Both OtUBD and 3xOtUBD bound (B) ubiquitylated substrates in the lysates. Concentration of bait protein indicates the amount of bait protein per unit volume of amylose resin. Bands with molecular sizes matching those of the MBP-OtUBD fusion proteins were also detected, potentially due to cross-reactivity of the ubiquitin antibody with OtUBD and the relatively large amounts of bait proteins present in the eluates. U: unbound fraction; B: bound fraction.

## Results

### OtUBD can protect and enrich ubiquitylated species from whole cell lysates

We first expressed and purified recombinant OtUBD with an N-terminal His_6_ tag (Fig. 1B). A previously reported TUBE based on the UBA domain of human UBQLN1 (4xTR-TUBE) was used for comparison (22, 43). One use of TUBEs is to protect ubiquitylated proteins *in vitro* from being cleaved by endogenous DUBs or being degraded by the proteasome following cell lysis, which facilitates their analysis (32). We tested if OtUBD could do the same. When yeast cells were lysed in the presence of NEM (N-ethylmaleimide, a covalent cysteine modifier that inhibits most cellular DUBs), 3 μM OtUBD or 3 μM TUBE, higher mass ubiquitylated species were similarly preserved by the two ubiquitin binders, with NEM having the strongest effect, as expected (Fig. 1C).

We investigated whether this protection extended to monoubiquitylated proteins by examining Flag-tagged histone H2B (Htb2) in a *ubp8Δ* mutant (44). Histone H2B is known to be monoubiquitylated, and levels of this species are enhanced by deleting Ubp8, the DUB that reverses the modification (45). Strikingly, OtUBD added to the cell lysate preserved the monoubiquitylated H2B to a degree comparable to NEM (Fig.1C, bottom). By contrast, H2B-ubiquitin was completely lost in extracts without any DUB inhibitor or when incubated with the TUBE protein.

We next determined if OtUBD or tandem repeats of OtUBD could be used for affinity enrichment of ubiquitylated proteins. We fused maltose-binding protein (MBP) to the N-terminus of OtUBD or three tandem OtUBD repeats (Fig. 1B). Purified MBP or the MBP fusion proteins were first bound to an amylose resin and then incubated with yeast whole cell lysates. With lower amounts of the resin-bound bait proteins, MBP-3xOtUBD enriched ubiquitylated proteins more efficiently than did MBP-OtUBD, likely due to increased avidity (Fig. 1D, left; compare bound (B) to unbound (U) lanes). When we increased the amount of the bait proteins, however, both MBP-OtUBD and MBP-3xOtUBD efficiently depleted ubiquitylated proteins from the lysate (Fig. 1D, right). The negative control MBP did not detectably bind any ubiquitylated species at either concentration. Notably, efficient enrichment was only achieved when MBP-OtUBD was pre-bound to the amylose resin (Fig. S1A). When free MBP-OtUBD was first incubated with the cell lysate and then bound to amylose resin, the enrichment efficiency was compromised (Fig. S1B). MBP-OtUBD also efficiently enriched ubiquitylated proteins from mammalian cell lysates, demonstrating its general utility across species (Fig. S1C).

In summary, OtUBD can both protect ubiquitylated proteins from *in vitro* deubiquitylation and enrich for such proteins. Unlike previously reported UBDs (32, 46), OtUBD can efficiently enrich ubiquitylated proteins even when used as a single entity instead of tandem repeats.

### A covalently linked OtUBD resin for ubiquitylated protein purification

We next generated resins with covalently attached OtUBD to minimize the contamination by bait proteins seen with MBP-OtUBD and maltose elution. Since OtUBD lacks cysteine residues, we introduced a cysteine residue at the N-terminus of the OtUBD sequence as a functional handle that can react with the commercially available SulfoLink™ resin to form a stable thioether linkage (Fig. 2A). As a negative control, free cysteine was added to the SulfoLink™ resin to cap the reactive iodoacetyl groups. When applied to yeast whole cell lysates prepared in a buffer with 300 mM NaCl and 0.5% Triton-X100 detergent, the OtUBD resin bound a broad range of ubiquitylated proteins and the bound proteins could be eluted with a low pH buffer (Fig. 2B, Fig. S1D; see Material and Methods). No ubiquitylated species were detected in the eluates from the control resin (Fig. 2B, Fig. S1D).

**Figure 2.**
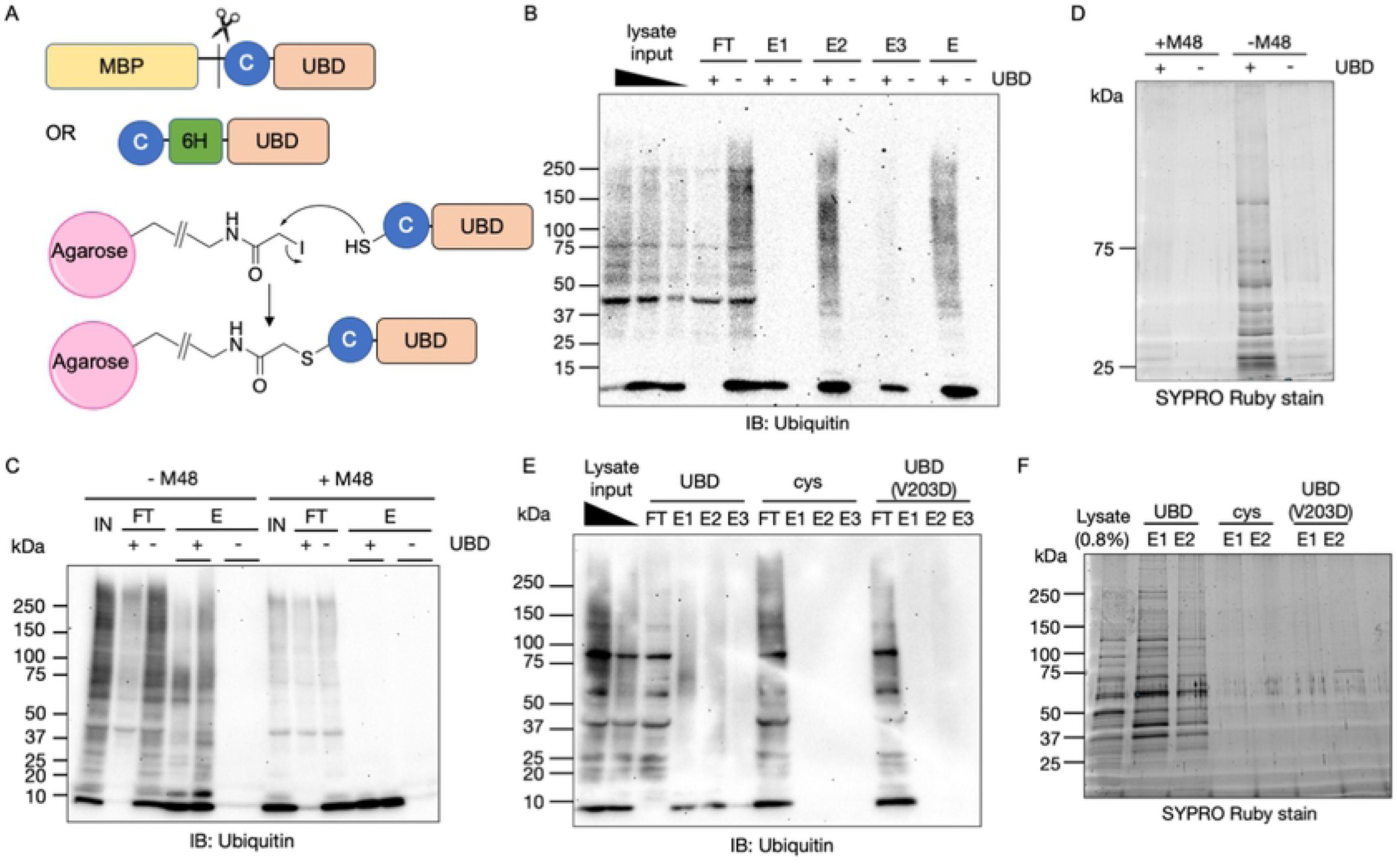
A covalently-linked OtUBD resin purifies ubiquitin and ubiquitylated proteins from yeast lysates. A. OtUBD constructs used for covalent coupling to resin and mechanism of the coupling reaction. An engineered cysteine at the N-terminus of OtUBD enables its covalent conjugation to the SulfoLink™ resin.
B. Ubiquitin blot of pulldowns from yeast cell lysate using covalently linked OtUBD resin or control resin. Covalently linked OtUBD resin efficiently pull down ubiquitylated species from yeast whole cell lysate. FT: flow-through; E1/E2/E3: eluted fractions using a series of stepwise, low pH elutions; E: pooled eluted fractions.
C, D. Extract pre-treatment with M48 DUB cleaves ubiquitin from ubiquitylated species and greatly reduces the total protein pulled down by OtUBD resin. C: Anti-ubiquitin blot of OtUBD pulldown of yeast lysate with or without M48 DUB treatment. D: Total protein present in the eluted fractions of the OtUBD pulldowns visualized with Sypro Ruby stain. (C and D are from two separate biological replicates.) IN: input; FT: flowthrough; E: eluted fractions.
E, F. The V203D mutation in OtUBD, which greatly impairs its binding of ubiquitin, prevents enrichment for ubiquitylated species from yeast lysate. E: Anti-ubiquitin blot of pulldowns of yeast lysates using OtUBD resin, Cys resin (negative control) and OtUBD(V203D) resin. F: Total protein present in the eluted fractions of the OtUBD pulldowns visualized with Sypro Ruby stain. IN: input; FT: flowthrough; E1/2/3: eluted fractions using a series of low pH elutions.

By comparing the anti-ubiquitin blot in Fig. 2B to the general protein stain of the same eluted fractions in Fig. S1D, it is clear that many proteins eluted from the OtUBD resin were not themselves ubiquitylated. Pulldown experiments performed under native or near-native conditions are expected to copurify proteins that interact noncovalently with ubiquitylated polypeptides, e.g., complexes that harbor ubiquitylated subunits. To test whether the entire protein population eluted from OtUBD resin was nevertheless dependent on substrate ubiquitylation, yeast lysates were pre-incubated with the viral M48 DUB, which cleaves a broad range of ubiquitylated proteins and reduces ubiquitin chains to free ubiquitin (Fig. 2C) (47). This treatment greatly reduced the total protein eluted from OtUBD resin compared to the pulldown from untreated lysate (Fig. 2D), indicating that the majority of proteins eluted from OtUBD resin were either ubiquitylated themselves or interacting with ubiquitin or ubiquitylated proteins.

A valine-to-aspartate mutation in OtUBD (V203D) severely impairs its binding to ubiquitin (41). To further validate the specificity of the OtUBD resin towards ubiquitylated proteins, we made an OtUBD(V203D) affinity resin and tested its ability to purify ubiquitin and ubiquitylated proteins. This single mutation greatly diminished the resin’s ability to enrich ubiquitylated species (Fig. 2E) and also strongly reduced the total bound protein eluate from the resin (Fig. 2F). This indicates that the ability of OtUBD resin to enrich for ubiquitylated species is based on its binding affinity towards ubiquitin.

Taken together, these results indicate the OtUBD resin specifically enriches ubiquitin and ubiquitylated polypeptides as well as proteins that interact with ubiquitin-containing proteins.

### Purifications using OtUBD with denatured extracts enrich ubiquitin-protein conjugates

To distinguish proteins covalently modified by ubiquitin from proteins co-purifying through noncovalent interaction with ubiquitin or ubiquitylated proteins, we optimized pulldown conditions to include a denaturation step (Fig. 3A). Yeast lysates were incubated with 8 M urea, a condition where the majority of proteins are unfolded, to dissociate protein complexes (48).

**Figure 3.**
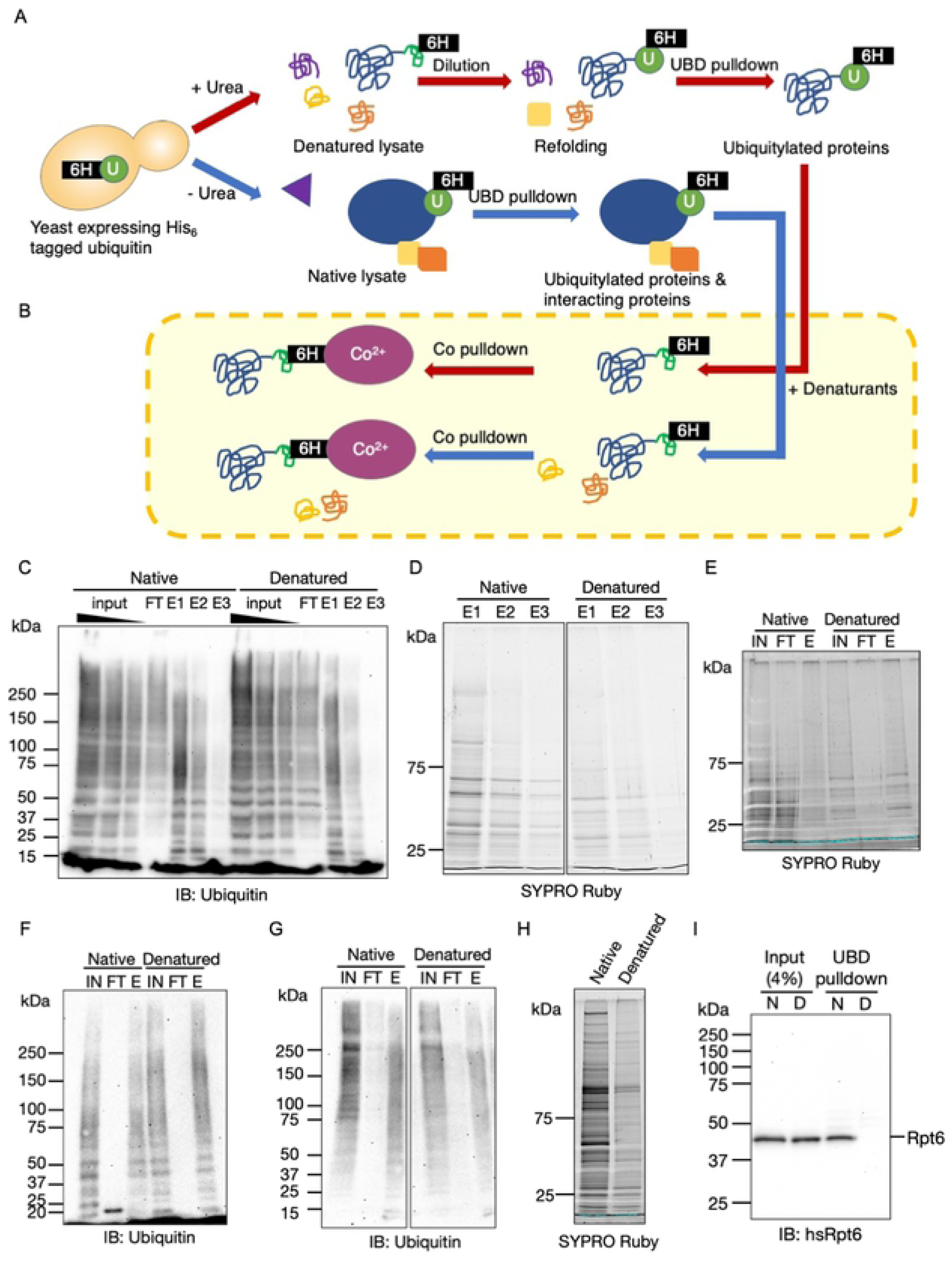
OtUBD pulldown under denaturing condition specifically enriches for proteins covalently modified with ubiquitin. A. Workflow of OtUBD pulldowns following sample denaturation (red arrows) or under native (blue arrows) conditions. In the first case, cell lysate is treated with 8 M urea to denature and dissociate proteins. The denatured lysate is then diluted 1:1 with native buffer to allow ubiquitin to refold and bind to OtUBD resin. Under such conditions, only ubiquitylated proteins are expected to be enriched. In the second case, cell lysate contains native ubiquitylated proteins as well as proteins that interact with them. OtUBD pulldown under such conditions is expected to yield both ubiquitylated substrates and ubiquitin-binding proteins.
B. Outline for the use of tandem Co^2+^ resin pulldowns to validate OtUBD pulldown results under different conditions. Eluates from OtUBD after lysates were incubated with denaturant (red arrows) or left untreated (blue arrows) are (re)treated with denaturant (8 M urea or 6 M guanidine•HCl) and then subjected to IMAC with a Co^2+^ resin in denaturing conditions. Proteins covalently modified by His_6_-ubiquitin bind to the Co^2+^ resin while proteins that only interact noncovalently with ubiquitin end up in the flowthrough.
C. Anti-ubiquitin blot of OtUBD pulldowns following native and urea denaturing treatments performed as described in Fig. 3A. FT: flowthrough; E1/2/3: eluted fractions from a series of low pH elutions.
D. Total protein present in eluates of the OtUBD pulldowns in Fig. 3C visualized by SYPRO Ruby stain. The image was spliced to remove irrelevant lanes.
E. Total protein present in different fractions of the Co^2+^ IMAC in Fig. 3E visualized by SYPRO Ruby stain.
F. Anti-ubiquitin blot of fractions from Co^2+^ IMAC (see Fig. 3B, the blot shown here used urea as the denaturant) of eluates from native and denaturing OtUBD resin pulldowns. IN: input; FT: flowthrough; E: fraction eluted with 500 mM imidazole.
G. Anti-ubiquitin blot of OtUBD pulldowns from HeLa cell lysates performed as described in Fig. 3A following native or denaturing treatments. The image was spliced to remove irrelevant lanes. IN: input; FT: flowthrough; E: fraction eluted with low pH elution buffer.
H. Total protein present in eluates in Fig. 3G visualized by SYPRO Ruby stain.
I. Immunoblot analysis of human proteasomal subunit Rpt6 in OtUBD pulldowns following native and urea denaturing treatments of lysates. Unmodified Rpt6 co-purified with OtUBD resin under native conditions but not following denaturation of extract. N: native condition; D: denaturing condition.

Denatured lysates were then diluted with native lysis buffer (to a final urea concentration of 4M) to facilitate the refolding of ubiquitin and applied to the OtUBD resin. A similar method was also used previously in ubiquitin immunoprecipitation using the FK2 monoclonal antibody (20).

Under such conditions, the OtUBD resin concentrated ubiquitylated proteins with efficiencies similar to those seen under native conditions (Fig. 3C). At the same time, the denaturing treatment greatly reduced the total amount of proteins eluted compared to native conditions, and the spectrum of purified protein species also changed (Fig. 3D). This suggests that ubiquitylated proteins were specifically enriched by the urea treatment.

To verify that OtUBD pulldown following a denaturation step is specific for proteins covalently modified with ubiquitin, we utilized a yeast strain whose endogenous ubiquitin-coding sequences were all deleted and replaced with a single plasmid-borne His_6_-tagged ubiquitin sequence (27). The eluted fractions from OtUBD resin pulldowns done after either denaturing or nondenaturing treatments of lysates (Fig. 3A) were then denatured again by incubation with urea or guanidine-HCl (Fig. 3B). The denatured proteins were then applied to a Co^2+^ (Talon) resin for immobilized metal affinity chromatography (IMAC) via the His_6_-tagged ubiquitin. If the eluate from the OtUBD resin had contained only (His_6_-)ubiquitylated proteins, most or all of the total proteins should bind to the resin. We observed that when OtUBD pulldowns were done following a denaturing lysate treatment, most of the eluted proteins were indeed bound to the Co^2+^ resin (Fig. 3E). By contrast, a large portion of proteins from a “native” OtUBD pulldown remained in the flow-through of the Co^2+^ resin (Fig. 3E). The overall levels of ubiquitylated species recovered, however, were comparable between the two treatments (Fig. 3F). Consistent with these findings with bulk ubiquitin conjugates, when we tested whether the proteasome, which binds noncovalently to many polyubiquitylated substrates (49), was in the OtUBD eluates, we readily detected proteasome subunits in the native pulldowns but not pulldowns from denatured lysates (Fig. S2A).

OtUBD-based affinity purifications, under either native or denaturing conditions, were also effective with human cell lysates. Both conditions led to similarly enrichment of ubiquitin conjugates (Fig. 3G), but the denaturing pretreatment greatly reduced the amounts of co-purifying nonubiquitylated proteins (Fig. 3H). Congruent with this, nonubiquitylated human proteasomal subunits were only present at substantial levels in eluates from native lysates (Fig. 3I, Fig. S2B). Interestingly, low amounts of presumptive ubiquitylated proteasome subunits were discovered in OtUBD pulldowns from both native and denatured lysates, and these species were strongly enriched over the unmodified subunits under the latter condition (Fig. S2B).

Overall, these results indicate that OtUBD-based protein purification under denaturing conditions can specifically enrich proteins that are covalently modified by ubiquitin.

### Enrichment using OtUBD to study ubiquitylation of specific proteins

One direct application of the OtUBD affinity reagent would be to facilitate the detection of ubiquitylated forms of specific proteins of interest. For example, histone H2B is monoubiquitylated by the ubiquitin E3 ligase Bre1 and deubiquitylated by the DUB Ubp8 in yeast (45, 50). The monoubiquitylated species of histone H2B is difficult to detect directly in the whole cell lysate due to its low abundance in comparison to unmodified H2B (Fig. 4A). To determine if the OtUBD resin could aid in the detection of monoubiquitylated H2B, we used OtUBD resin to purify total ubiquitylated proteins from cell lysates of wild-type (WT), *bre1Δ* and *ubp8Δ* yeast strains expressing Flag-tagged histone H2B and then analyzed the proteins by immunoblotting. A slower migrating band in the anti-Flag immunoblot, which represents the monoubiquitylated H2B, was detected in the WT and *ubp8Δ* yeasts but not in the *bre1Δ* yeast (Fig. 4A). By contrast, the 4xTR-TUBE-resin failed to capture the monoubiquitylated H2B in any of the yeast samples (Fig. 4A), including those with elevated levels of H2B monoubiquitylation due to deletion of *UBP8* (Fig. 4A). Thus, although 4xTR-TUBE-resin can efficiently enrich bulk ubiquitylated species from these lysates (Fig. S3A), this approach can be limited in its ability to detect monoubiquitylated proteins. These results again highlight a potential advantage of OtUBD over TUBEs in studying monoubiquitylated substrates.

**Figure 4.**
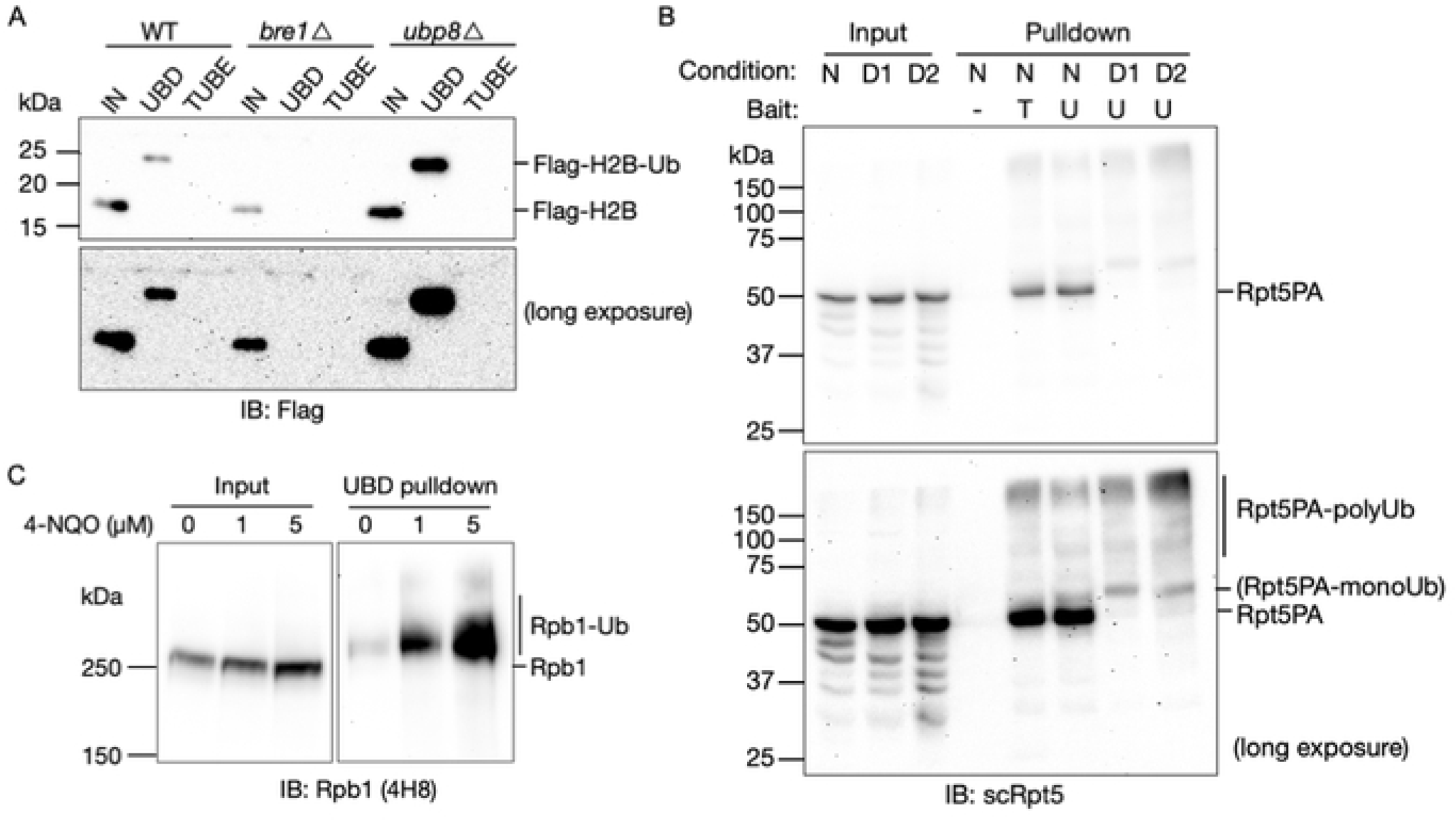
OtUBD resin as an enrichment tool to study ubiquitylation of specific proteins. A. Anti-Flag immunoblot of OtUBD and TUBE pulldowns from lysates of WT, *bre1Δ* and *ubp8Δ* yeasts expressing Flag-tagged histone H2B. OtUBD resin but not TUBE bound monoubiquitylated histone H2B from whole cell lysate of WT and *ubp8Δ* yeasts. IN: input; UBD: eluted fraction from OtUBD pulldown; TUBE: eluted fraction from TUBE pulldown. Elution was achieved by incubating resins with SDS sample buffer.
B. Western blot of yeast Rpt5 from pulldowns of *rpt2-P103A rpt5-P76A* (*rpt2,5PA*) mutant yeast lysates using different ubiquitin affinity resins under different conditions. N: pulldown performed under native conditions; D1: pulldown performed after 8 M urea denaturation as described in Fig. 3A; D2: pulldown performed after urea denaturation directly in buffer containing 8 M urea. −: negative control resin; T: TUBE resin; U: OtUBD resin.
C. Western blot analysis of RNAPII subunit Rpb1 in OtUBD pulldowns of denatured HeLa cell lysates. Cells were treated with increasing concentrations of 4-NQO to induce RNAPII ubiquitylation. The image was spliced to remove an empty lane for presentation.

Our lab previously identified mutations in yeast proteasomal subunits Rpt2 and Rpt5 (*rpt2,5PA*) that lead to their misfolding and ubiquitylation under normal growth conditions (51). In that study, the ubiquitylation of the Rpt subunits was confirmed by overexpressing His-tagged ubiquitin in the proteasome mutant yeast strain and performing IMAC under denaturing conditions to capture ubiquitylated species. We performed OtUBD and TUBE pulldowns with *rpt2,5PA* yeast lysates without ubiquitin overexpression. Based on anti-Rpt5 immunoblotting, both resins captured a smear of higher mass Rpt5PA species, which are likely endogenous polyubiquitylated Rpt5PA species (Fig. 4B). Unmodified Rpt5PA co-purified with both OtUBD and TUBE under native conditions but was largely eliminated from OtUBD pulldowns done after lysate denaturation. Compared to the TUBE pulldown, OtUBD pulldown captured an additional lower molecular weight Rpt5PA species which, based on the apparent molecular mass, is likely monoubiquitylated Rpt5PA (Fig. 4B).

As a final example of single protein analysis, we used OtUBD to detect ubiquitylated RNA polymerase II (RNAPII) in cultured human cells. RNAPII becomes ubiquitylated upon UV-induced DNA damage (52). Rpb1, the largest subunit of RNAPII, is heavily ubiquitylated under such conditions (53). We treated HeLa cells with the chemical 4-NQO (4-nitroquinoline-1-oxide), which mimics the biological effects of UV on DNA (54), and performed OtUBD pulldowns of both native (Fig. S3B) and denatured (Fig. 4C) lysates. In both cases, similar slower migrating bands were present in the eluted fractions analyzed by anti-Rpb1 immunoblotting. Because Rpb1 is a large protein of over 200 kDa and exists in different phosphorylation states, it is difficult to distinguish nonubiquitylated and monoubiquitylated species based on migration through an SDS-PAGE gel (55). OtUBD pulldown under denaturing conditions, in this case, provides confidence that Rpb1 is ubiquitylated even under basal conditions and becomes heavily ubiquitylated upon treatment with 4-NQO (Fig. 4C).

These examples illustrate how OtUBD resin can facilitate the detection of monoubiquitylated and polyubiquitylated proteins of interest in both yeast and human cells.

### OtUBD-pulldown proteomic profiling of the yeast and human ubiquitylome and ubiquitin interactome

By comparing OtUBD pulldowns of native and denatured cell lysates, we can potentially differentiate different ubiquitin-related proteomes in a biological sample. The “ubiquitylome,” i.e., the collection of covalently ubiquitylated proteins, can be defined as the protein population eluted from an OtUBD affinity resin used with denatured cell extracts. The “ubiquitin interactome” can be roughly defined as those proteins that are specifically enriched following OtUBD pulldowns from native extracts but not pulldowns from denatured lysates (Fig. 4A).

Notably, the latter definition will exclude cases where a subpopulation of a protein is ubiquitylated while the non-ubiquitylated population of the same protein interacts noncovalently with ubiquitin or other ubiquitylated proteins. For example, some proteasomal subunits are known to be ubiquitylated (56), but proteasome particles where these subunits are unmodified still interact noncovalently with ubiquitylated proteins. Proteins such as these proteasomal subunits will be excluded from the ubiquitin interactome as defined here. Nevertheless, these definitions provide a general picture of the ubiquitylome and ubiquitin interactome.

We performed OtUBD pulldowns of whole yeast lysates with and without prior denaturation (Fig. S4A-C) and profiled the eluates using shotgun proteomics. For each condition, we included two biological replicates and for each biological replicate, two technical repeats of the LC-MS/MS runs. Control pulldowns by SulfoLink™ resin without OtUBD were performed in parallel to eliminate proteins that non-specifically bind to the resin. As was seen in earlier experiments, the control pulldowns yielded no detectable ubiquitylated species and at most trace amounts of proteins (Fig. S4A-C). Some proteins were identified in a subset of the control pulldown replicates (Fig. S4D), due partially to carryover of high abundance peptides from previous runs, but the overall quantities of proteins in these control samples, as demonstrated by total TIC (total ion current), were much lower compared to the OtUBD pulldown samples (Fig. S4E). Hence, for each biological replicate, only proteins present at significantly higher levels (>20 fold) in the OtUBD pulldown samples over the corresponding control pulldown samples were considered real hits (Fig. S4F, Supplementary Data 2).

The two pulldown conditions yielded similar total numbers of proteins (Fig. 5A) with a major overlap of protein identities. Over 400 proteins were discovered exclusively under native conditions, suggesting they are not ubiquitylated themselves but co-purify with ubiquitin or ubiquitylated proteins. Interestingly, over 600 proteins were identified only under denaturing conditions. Because OtUBD pulldowns following lysate denaturation yield much less total protein than under native conditions (Fig. S4C), the possibility of identifying low abundance proteins in the LC-MS/MS analysis is likely increased.

**Figure 5.**
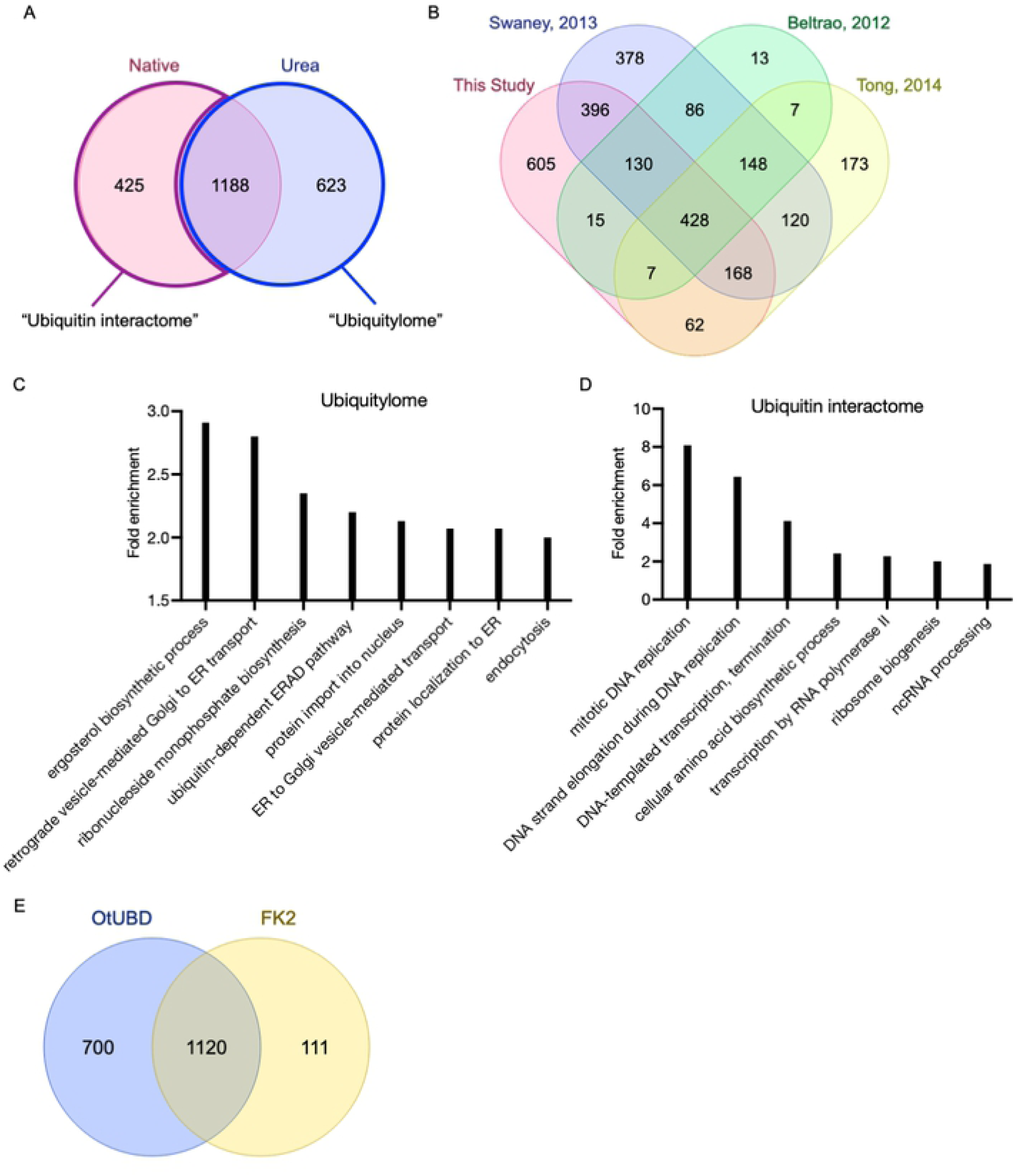
OtUBD pulldown-proteomics enables profiling of the ubiquitylome and ubiquitin interactome of yeast and human cells. A. Venn diagram of yeast proteins identified by OtUBD pulldown-proteomics with the pulldowns performed under following either nondenaturing or urea denaturing treatments. The collection of proteins identified in OtUBD pulldowns under denaturing condition is defined as the ubiquitylome (blue outline). The collection of proteins identified only in native OtUBD pulldown is defined as the ubiquitin interactome (purple outline).
B. Venn diagram comparing the yeast ubiquitylome defined by OtUBD pulldown-proteomics with three previous studies using the di-Gly antibody IP method (57–59).
C, D. Top biological pathways involved in OtUBD pulldown-defined yeast ubiquitylome (C) and ubiquitin interactome (D) based on GO analysis.
E. Venn diagram comparing the human ubiquitylome defined by OtUBD pulldown and FK2 antibody IP.

We compared the OtUBD-defined yeast ubiquitylome with data from previously published studies using di-Gly remnant antibody-based methods (Fig. 5B) (57–59). Our study identified 1811 ubiquitylated yeast proteins, the second highest number among the four studies compared here. About two-thirds of proteins identified in our study have been reported to be ubiquitylated by at least one of these di-Gly antibody-based studies. Around 600 ubiquitylated proteins were uniquely identified in this study. Some of these might involve non-canonical ubiquitylation, where the ubiquitin modifier is covalently attached to a nucleophilic residue on the substrate other than a lysine (3).

GO analysis indicated that the yeast ubiquitylome defined by OtDUB binding spans proteins from a wide variety of cellular processes, including multiple biosynthesis pathways, protein localization, vesicle-mediated transport, and protein quality control pathways (Fig. 5C). By contrast, the ubiquitin interactome, as defined above, appeared to yield greater representation in nucleic acid-related processes such DNA replication, RNA transcription, ribosome biogenesis and noncoding RNA processing (Fig. 5D).

We also performed OtUBD pulldowns with denatured HeLa cell lysates and compare the data side by side with results obtained from immunoprecipitation using the FK2 antibody, a monoclonal antibody raised against ubiquitin (20). Both OtUBD and FK2 antibody resins efficiently enriched ubiquitylated proteins from HeLa cell lysates (Fig. S4D). The majority of ubiquitylated proteins identified by the FK2 antibody resin were also found in the ubiquitylated proteome identified by OtUBD resin (Fig. 5E). Compared to the FK2 immunoprecipitation, OtUBD pulldowns identified 700 additional ubiquitylated proteins, the majority of which have at least one reported ubiquitylation site in a previous study using diGly antibodies (60).

These proteomics experiments demonstrated that the OtUBD affinity resin can be used to profile the ubiquitylated proteome of both yeast and human cells. Moreover, all seven lysine-linkages of polyubiquitin chains were identified in the yeast proteomics (Fig. S4E) and their ratio roughly agree with previous quantitative studies of relative linkage frequencies (4). All except the K33 ubiquitin-ubiquitin linkage was also identified in the HeLa proteomics analysis. We note that K33 is a low abundance linkage (4), and only one biological replicate was analyzed for the HeLa cell experiment.

### OtUBD and label-free quantitation enable identification of potential E3 ligase substrates

Finally, we sought to apply OtUBD-pulldown proteomics toward identifying substrates of specific E3 ligases. Identification of substrates for particular E3 ligases can be challenging due to the transient nature of E3-substrate interaction and the low abundance and instability of many ubiquitylated proteins (61). One way to screen for potential substrates is to compare the ubiquitylome of cells with and without (or with reduced level/function of) the E3 of interest (62). Proteins with higher ubiquitylation levels in the cells expressing the E3 compared to cells lacking it would be candidate substrates. We used OtUBD-pulldown proteomics to profile the ubiquitylomes of wild type BY4741 yeast and two congenic yeast E3 deletion strains obtained from a gene knockout library (63). Of the E3s we chose to study, Bre1 is a relatively well-characterized ligase that monoubiquitylates histone H2B (Fig. 4A) (50). This ubiquitylation does not lead to H2B proteolysis but is involved in important cellular processes including transcription and DNA damage repair (64). Other substrates of Bre1 are largely unknown. The other E3 Pib1 is a much less studied ligase that localizes to the endosomes and the vacuole and participates in endosomal sorting (65).

We harvested WT, *bre1Δ* and *pib1Δ* yeast cells and performed OtUBD pulldowns following lysate denaturation. Proteins eluted from the OtUBD resin were subject to label-free quantitative proteomics (Fig. 6A, Fig. S5A, B). Three biological replicates were examined for each group, and each replicate was analyzed by two separate LC-MS/MS runs. Quantitation was performed using total TIC (total ion current) after normalization among the analyzed samples. As expected, histone H2B (identified as Htb2) presented at a much higher level in the ubiquitylome of WT cells compared to that of *bre1Δ* cells (Fig. 6B). Interestingly, we identified two different ubiquitylation sites on histone H2B (Htb2) in different samples (Fig. 6C, Fig. S6A, B). The K123 ubiquitylation site, which is the major reported ubiquitylation site of Bre1 on histone H2B (66, 67), only showed up in WT cells but not in *bre1Δ* cells. By contrast, the other ubiquitylation site, K111, showed up in both WT and *bre1Δ* cells. This indicates that there is an E3 ligase(s) other than Bre1 that can ubiquitylate histone H2B on K111. Although this site had been reported in a diGly antibody-based proteomics study (58), the function of this ubiquitylation remained to be studied. Besides histone H2B, we also identified 16 other proteins present in significantly higher levels in the WT cell ubiquitylome compared to *bre1Δ* cells (Fig. 6D). In addition, 35 proteins were only detected in the ubiquitylome of WT cells but not *bre1Δ* cells (Fig. 6E). Taken together, these proteins are considered potential Bre1 substrates. Interestingly, some of these proteins (Fig. 6D, E, green) have been shown to be metabolically stabilized in *bre1Δ* cells in an earlier study (68), which indicates that they could be direct or indirect proteolytic ubiquitylation substrates of Bre1.

**Figure 6.**
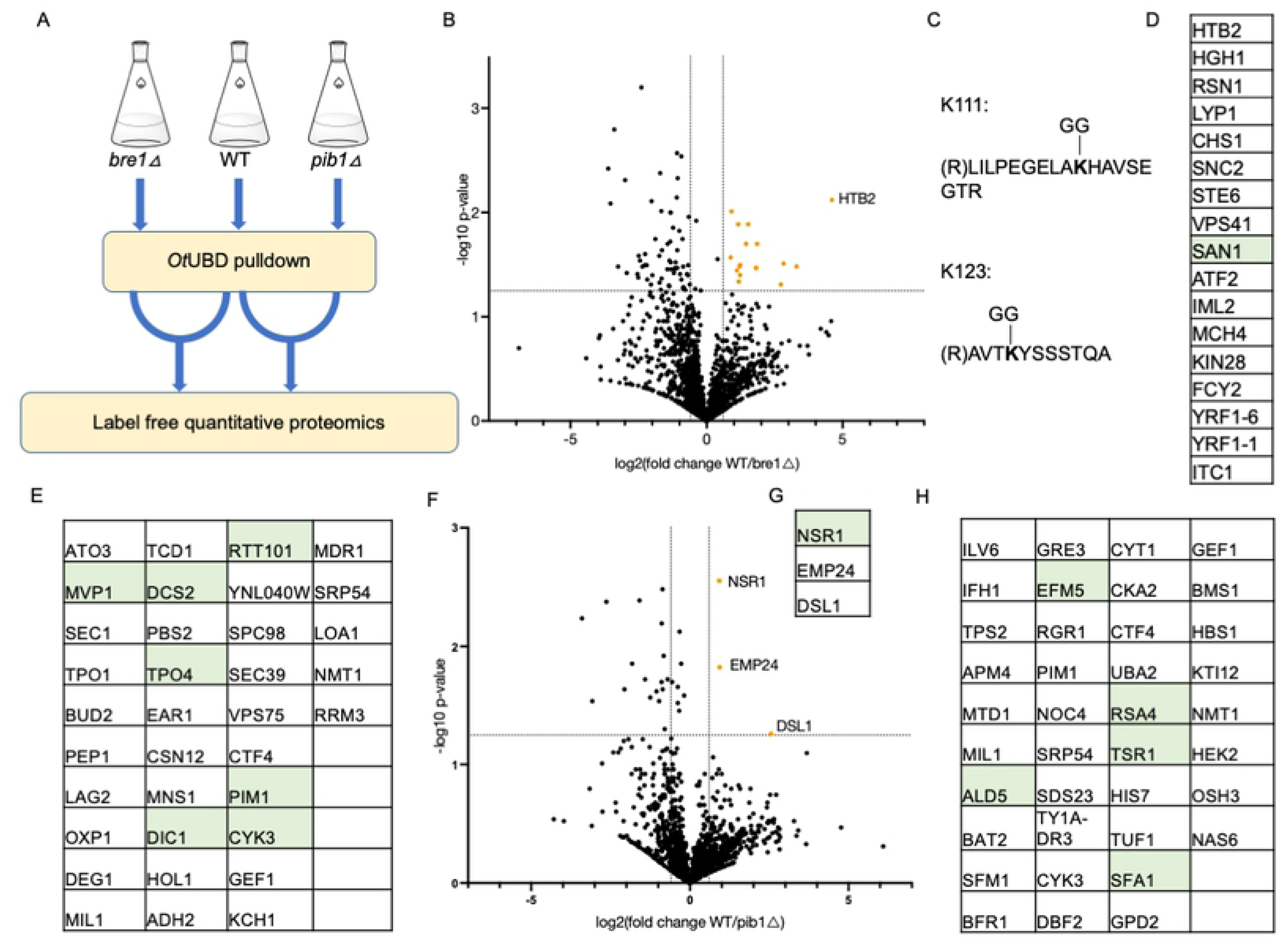
Identification of potential E3 substrates by OtUBD pulldown and label-free quantitation. A. Scheme for E3 substrate identification using OtUBD pulldown and quantitative proteomics. WT and E3 deletion (*bre1Δ* and *pib1Δ*) yeast strains were subjected to OtUBD pulldowns following extract denaturation. The eluted proteins were then analyzed by label-free quantitation.
B. Volcano plot comparing WT and *bre1Δ* samples. Orange dots represent proteins that were significantly enriched in WT samples compared to *bre1Δ* samples. Horizontal dashed line indicates p = 0.05. Vertical dash lines indicate relative change of +/- 1.5-fold.
C. Two different ubiquitylation sites identified on histone H2B (Htb2) in different samples.
D. List of proteins that were significantly enriched in WT samples compared to *bre1Δ* samples (orange dots in B). Green color indicates proteins that were previously reported to be stabilized in *bre1Δ* yeast.
E. Ubiquitylated proteins detected exclusively in WT but not *bre1Δ* samples. Green color indicates proteins previously reported to be stabilized in *bre1Δ* cells.
F. Volcano plot comparing WT and *pib1Δ* samples. Orange dots represent proteins that were significantly enriched in WT samples compared to *pib1Δ* samples. Horizontal dash line indicates p = 0.05. Vertical dash lines indicate relative change of +/- 1.5-fold.
G. List of proteins that were significantly enriched in WT samples compared to *pib1Δ* samples (orange dots in F). Green color indicates proteins that were previously reported to be stabilized in *pib1Δ* yeast.
H. Ubiquitylated proteins detected exclusively in WT but not *pib1Δ* samples. Green color indicates proteins previously reported to be stabilized in *pib1Δ* yeast.

Analogous to the Bre1 data, we identified three proteins whose ubiquitylated forms were found at significantly higher levels in WT cells versus *pib1Δ* cells (Fig. 6F, G) and 38 proteins that were detectably ubiquitylated only in WT cells but not *pib1Δ* cells (Fig. 6H). Of these proteins, six have been shown previously to be stabilized in *pib1Δ* cells (Fig. 6G, H) (68).

Whether these potential E3 substrates are direct or indirect ubiquitylation substrates of the tested E3s will need to be validated by biochemical assays. Nonetheless, our results demonstrated that OtUBD can serve as a means to profile ubiquitylomes quantitatively, which could be useful in the identification of substrates for E3 ligases and other ubiquitin-related enzymes such as E2s and DUBs.

Among the various proteomics data obtained from our OtUBD pulldowns, we observed a number of potential non-lysine ubiquitylation sites assigned by the Mascot search algorithm (Supplementary Data 2), substantiating the idea that OtUBD can enrich proteins with non-lysine ubiquitylation sites. We confirmed one of these sites by manual validation of the spectrum assignment (Fig. S6C).

## Discussion

Protein ubiquitylation continues to be of great interest due its vital contributions to many fundamental cellular processes and for its important roles in human disease. Many enzymes involved in ubiquitylation are being pursued as targets for therapeutics (69, 70). For example, a number of drug candidates targeting E3 ligases such as MDM2 and XIAP have entered clinical trials for treatment of multiple types of cancer (71). A variety of reagents and methods to study ubiquitylation or ubiquitylation-related processes have been developed, but these methods all have limitations (16, 39). For example, TUBEs are effective at detecting polyubiquitin chains, but this creates a bias towards polyubiquitylated substrates; they often fail to detect protein monoubiquitylation signals (e.g., Fig. 4A), which can dominate the ubiquitylome in at least some mammalian cell types (17). As a result, new and economical reagents and methods to analyze the many types of ubiquitin modification are still needed, particularly when these modifications are present at very low levels.

The versatile high-affinity UBD domain of *O*. *tsutsugamushi* DUB provides an affinity reagent with several advantages over existing tools. First, it is straightforward and relatively inexpensive to generate the affinity resin using the small recombinant OtUBD protein expressed and purified from *E. coli*. Second, ubiquitin enrichment using OtUBD is applicable to both monoubiquitylation and polyubiquitylation, in contrast to the bias of TUBEs and other reagents that depend on binding avidity for binding polyubiquitin. Third, OtUBD pulldowns can be performed under native conditions for the study of both ubiquitylated substrates and proteins that associate noncovalently with them, or by subjecting extracts to denaturing conditions prior to pulldown, OtUBD pulldowns can be tuned toward proteins covalently modified by ubiquitin.

OtUBD pulldowns, coupled with proteomics, can be used to profile the ubiquitylated proteomes of yeast and mammalian cells and no doubt other eukaryotic cells. Fourth, when done quantitatively, comparative OtUBD pulldown-proteomics can be used to identify substrates of ubiquitylating enzymes (E2s or E3s), as shown here, or DUBs (unpublished results). Finally, unlike the anti-diGly immunoaffinity tool that is specific for diGly remnants on Lys side chains, OtUBD-based purifications can help identify noncanonical ubiquitin-protein linkages such as through Cys, Ser, or Thr side chains, the N-terminal amino group, or protein bonds that do not involve the ubiquitin C-terminus, as in ubiquitylation mediated by *Legionella* SidE proteins (3, 72, 73). It should also be possible to enrich monoubiquitin linkages to macromolecules other than proteins, such as the recently discovered ubiquitin-lipopolysaccharide adducts formed during *Salmonella* infections (40).

We have demonstrated that OtUBD is specific towards ubiquitin and ubiquitylated proteins. However, several caveats should be noted. Although OtUBD pulldowns following extract denaturation significantly reduces the amount of interacting proteins co-purifying with ubiquitylated proteins, a small number of noncovalently interacting proteins may still be co-purified in some cases (e.g. Fig. S2B). Additional stringent wash steps may help mitigate this problem. OtUBD also binds to the closely related ubiquitin-like modifier Nedd8, although with a much lower affinity than for ubiquitin (41). Like ubiquitin, Nedd8 is used for protein post-translational modification (74) and because they leave a same −GlyGly remnant residue after trypsin digestion, it is hard to differentiate the two modifiers using the normal diGly antibody method (16). We looked for potential neddylation substrate(s) in our proteomics studies. Rub1 (yeast Nedd8), Cdc53, Rtt101 and Cul3 (three yeast cullin proteins reported to undergo neddylation (75)) were detected in the OtUBD-defined ubiquitylome, which may have been enriched on the OtUBD resin through Nedd8 binding. Nevertheless, neddylation occurs at much lower levels compared to ubiquitylation (37) and based on the specificity analysis we performed (Fig. 2D, Fig. 3E), neddylated proteins (if any) should account for only a small fraction of the OtUBD-enriched proteome.

In our OtUBD pulldown-proteomics experiments, the total number of ubiquitylated yeast proteins identified was comparable to previous studies using the di-Gly antibody enrichment method (57–59). Optimization of our proteomics pipeline would likely further improve results, especially for low-abundance proteins. In the samples we analyzed, peptides derived from ubiquitin accounted for a significant percentage of the total number of identified peptides. This likely limited detection of low-abundance peptides, especially those with similar retention time. Pre-clearing of free ubiquitin from the eluted samples before LC-MS/MS, for example, by gel separation or an affinity depletion specific for free ubiquitin (76), would likely reduce this problem. Additional fractionation of the protein or peptide samples should also enhance the overall discovery rate.

As a ubiquitin enrichment method on the protein level, our method could be used in conjunction with other methods for efficient enrichment of certain ubiquitylated species. For example, OtUBD pulldowns could be performed before di-Gly antibody immunoprecipitation (IP). When performed against the enormous pool of peptides derived from the entire cell proteome, di-Gly antibody IP often needs to be done in multiple batches or for multiple rounds to ensure efficient enrichment (60). A preliminary OtUBD pulldown step could significantly clean up the sample without creating any bias towards polyubiquitylated species. This will greatly increase the percentage of di-Gly-linked peptides present in the digested sample. Since OtUBD has exceptionally high affinity towards free ubiquitin, it could also be used with the Ub-Clipping technique (77). In Ub-Clipping, ubiquitylated proteins are cleaved at the ubiquitylation sites by the protease Lb-Pro to generate diGly-linked monoubiquitin species and free ubiquitin_1-74_. These species carry information on ubiquitin chain topology and post-translational modifications of ubiquitin that can be deciphered by MS analysis. Deployment of OtUBD for other applications can be readily envisioned.

With the characterization of OtUBD-ubiquitin binding and crystal structures of OtUBD available (41), one can imagine further modifications that would adapt or enhance OtUBD for other uses. For example, directed evolution or structure-based rational mutagenesis may be performed to change OtUBD binding specificity toward ubiquitin, specific ubiquitin chains or ubiquitin-like proteins. OtUBD could be turned into a ubiquitin detection tool for other applications by attaching a fluorophore or other functional handles to it. As a recombinant protein reagent that is versatile and easy to prepare, OtUBD will also be an economical addition to the ubiquitin research toolbox.

## Materials and Methods

### Plasmids and DNA cloning

The coding sequence for 3xOtUBD was synthesized by Genscript USA. pRSET-4xTR-TUBE was a gift from Yasushi Saeki (Addgene plasmid # 110312) (43). All DNA constructs made in this study were based on either the 4xTR-TUBE or OtUBD insert. The pRT498 vector, a bacterial expression plasmid modified from pET42b to include an N-terminal His_6_-MBP with a cleavable TEV site, was used for expression of MBP and MBP-fusion proteins made in our lab. pET21a and pET42b vectors were used to express His_6_-tagged proteins in bacteria. Plasmids and primers used in this study are described in detail in Supplementary Data 1. All PCR reactions were done using Phusion® High-Fidelity DNA Polymerase (New England Biolabs).

### Yeast strains and growth

Yeast strains used are listed in Supplementary Data 1. Yeast cultures were grown overnight in yeast extract-peptone-dextrose (YPD) medium to saturation. The next day, the culture was diluted in fresh YPD to an OD_600_ of 0.1 to 0.2, and cultured at 30°C with shaking until reaching mid-exponential phase (OD_600_ 0.8-1.2). Cells were pelleted, washed with water, and flash frozen in liquid nitrogen and stored at −80 °C until use.

### Mammalian cell culture

HeLa and HEK293T cells (ATCC) were cultured in Dulbecco’s Modified Eagle Medium (Gibco) supplemented with 10% fetal bovine serum (Gibco) and 1% penicillin-streptomycin (Gibco). Cells were not used past 20 passages. To harvest cells for experiments, the medium was removed, and cells were washed with cold Dulbecco’s phosphate-buffered saline (DPBS, Gibco) before dislodging by scraping in cold DPBS. The dislodged cells were pelleted by centrifugation at 400 x *g* for 4 minutes and flash frozen in liquid nitrogen. Cell pellets were stored at −80 °C until use.

### Expression and purification of recombinant proteins

Recombinant His_6_-tagged proteins were purified from Rosetta (DE3) competent *E. coli* cells (Novagen) transformed with the appropriate plasmids. Bacterial cells were grown overnight in Luria-Bertani (LB) broth supplemented with either 100 μg/mL ampicillin (for pET21a-based plasmids) or 50 μg/mL kanamycin (for pRT498- and pET42b-based plasmids) and diluted 1/100 the next morning in fresh LB broth supplemented with the corresponding antibiotics. When cell density had reached 0.5–0.8 OD_600_, protein production was induced by addition of isopropyl β-D-1-thiogalactopyranoside (IPTG) to a final concentration of 0.3 mM and cells were cultured at 16-18°C overnight with shaking. Bacteria were pelleted and resuspended in bacteria lysis buffer (50 mM Tris•HCl, pH 8.0, 300 mM NaCl, 10 mM imidazole, 2 mM phenylmethylsulfonyl fluoride (PMSF)) supplemented with lysozyme and DNaseI, incubated on ice for 30 minutes and lysed using a French press. Lysates were clarified by centrifugation for 1 h at 4°C at 10,000 rcf before being subjected to Ni-NTA (Qiagen) affinity purification following the manufacturer’s protocol.

For purification of OtUBD variants using pRT498-based plasmids, the proteins eluted from the Ni-NTA resin were subject to buffer exchange in a 50 mM Tris•HCl, pH 7.5, 150 mM NaCl buffer supplemented with 10 mM tris(2-carboxyethyl)phosphine) (TCEP) (from a 1M TCEP stock neutralized with NaOH to pH 7) using a centrifugal filter device (Amicon, 3000 MWCO) following manufacturer’s protocol. His-tagged TEV protease was added to remove the His_6_-MBP tag, and the mixture was incubated on ice overnight. The cleavage mixture was then allowed to flow through a column of clean Ni-NTA resin to capture the cleaved His_6_-MBP tag. The flow-through was concentrated and purified by Fast Protein Liquid Chromatography (FPLC) with a Superdex 75 (Cytiva) gel filtration column using FPLC buffer (50 mM Tris•HCl, pH 7.5, 150 mM NaCl, 1 mM TCEP). For further purification of His_6_-tagged OtUBD or 4xTR-TUBE, the protein eluate from the Ni-NTA matrix was supplemented with 5 mM TCEP, concentrated and purified by FPLC on a Superdex 75 gel filtration column using FPLC buffer containing 1 mM TCEP. For purification of MBP-tagged proteins, the protein eluate from the Ni-NTA resin was concentrated and fractionated by Superdex 75 FPLC using 50 mM Tris•HCl, pH 7.5, 150 mM NaCl buffer supplemented with 1 mM dithiothreitol (DTT). The M48 DUB protein was prepared as described earlier (47).

All proteins were flash-frozen in liquid nitrogen and stored at −80 °C until use. Protein concentrations were determined by either SDS-PAGE and GelCode Blue (Thermo) staining or a BCA assay (Thermo) using bovine serum albumin (BSA) as the standard.

### Immunoblotting and antibodies

Proteins resolved through SDS-PAGE gels were transferred to Immobilon-P PVDF membranes (Millipore) and blocked with 3% non-fat milk in Tris-buffered saline with 0.1% Tween-20 (TBST). Membranes were incubated first with the desired primary antibody diluted in TBST containing 1% milk for 1 hour at room temperature or overnight at 4°C, washed extensively, and then incubated with an HRP-linked secondary antibody diluted in TBST with 1% milk for 1 hour at room temperature or overnight at 4°C.

Primary antibodies used in this study were rabbit polyclonal anti-ubiquitin antibody (Dako, discontinued, 1:2000 dilution), monoclonal mouse anti-Flag M2 (Sigma, 1:5000 or 1:10000), monoclonal mouse anti-human Rpt6 (PSMC5) (Invitrogen, 2SU-1B8, 1:10000), monoclonal mouse anti-human Rpt4 (Enzo, p42-23, 1:1000), mouse monoclonal anti-yeast Rpt4 (gift from W. Tansey, 1:2500), rabbit polyclonal anti-Rpt5 (Enzo Life Sciences), rabbit polyclonal anti-Pre6 (gift from D. Wolf, 1:5000) and anti-Rpb1 (RNA Pol II) monoclonal mouse antibody (Active Motif, 4H8, 1:2000). For rabbit primary antibodies, the HRP-linked anti-rabbit IgG secondary antibody (GE Healthcare, NA934) was used at a dilution of 1:5000 or 1:10000. For mouse primary antibodies, the HRP-linked anti-mouse secondary antibody (GE Healthcare, NXA931V) was used at a dilution of 1:10000.

Blots were visualized by enhanced chemiluminescence on a G:Box imaging system with GeneSnap software (Syngene). Images were processed with ImageJ software.

### Protection of ubiquitylated species in whole cell yeast lysates

Yeast *ubp8*Δ cells expressing Flag-tagged histone H2B (44) were grown in YPD medium and harvested during exponential phase growth. The cell pellet was washed with water, flash-frozen, and lysed by grinding under liquid nitrogen in a mortar. Proteins were extracted by addition of lysis buffer (50 mM Tris•HCl, pH 7.5, 150 mL NaCl, 1 mM ethylenediaminetetraacetic acid (EDTA), 10% glycerol, cOmplete™ EDTA-free protease inhibitor cocktail (Roche), 1mM PSMF) in the presence of 20 mM N-ethylmaleimide (NEM), 3 μM OtUBD, 3 μM 4xTR-TUBE (all final concentrations), or nothing. The resulting lysates were cleared by centrifugation at 21,000 x *g* for 12 minutes at 4°C and incubated at room temperature for 1-4 hours. Flag-tagged H2B was purified by anti-Flag immunoprecipitation with ANTI-FLAG® M2 Affinity Gel (Millipore) following the manufacturer’s protocol. Whole cell lysates were analyzed by anti-ubiquitin immunoblotting. The anti-Flag precipitates were analyzed by anti-Flag immunoblotting.

### Pulldown with MBP-tagged bait proteins

Pulldowns of MBP-tagged fusion proteins were performed using an amylose resin (New England Biolabs). Appropriate amounts of (see Fig. 1D, S1A and S1B) MBP or MBP fusion proteins diluted in 300 μL amylose column buffer (20 mM Tris•HCl, pH 7.5, 200 mM NaCl, 1 mM EDTA, 1 mM DTT) were incubated with 50 μL amylose resin for 1 hour at 4°C with rotation. The resin was pelleted by centrifugation at 5,000 x *g* for 30 seconds, and the supernatant was removed. One mL of yeast lysate (1-2 mg/mL) prepared in column buffer freshly supplemented with protease and DUB inhibitors (cOmplete mini EDTA-free (Roche), 10 mM NEM, 2 mM PSMF) was added to the beads. (For detailed methods of lysate preparation, see section “Ubiquitin pulldown with protein-linked resins” below.) The mixture was incubated with rotation at 4°C for 2 hours. The resin was washed 5 times with 1 mL column buffer and then eluted by incubating with column buffer containing 50 mg/mL maltose for 2 hours at 4°C with rotation. Alternatively, bound proteins could be eluted by incubating with SDS sample buffer for 15 minutes at room temperature.

In the alternative incubation method described in Fig. S1B, MBP-OtUBD was first incubated with yeast lysate for 4 hours at 4°C with rotation. The mixture was then added to the amylose resin and incubated with rotation at 4°C for another 2 hours, followed by the same washing and elution steps described above.

### Generation of covalent-linked affinity purification resins

#### OtUBD resin

Covalently linked OtUBD resin was made by conjugating Cys-OtUBD or Cys-His_6_-OtUBD to SulfoLink™ coupling resin (Thermo Scientific) according to the manufacturer’s protocol. Briefly, 2 mL (bed volume) of SulfoLink resin was placed in a gravity column and equilibrated with 4 bed volumes of SulfoLink™ coupling buffer (50 mM Tris•HCl, 5 mM EDTA, pH 8.5). Four mg of Cys-OtUBD was diluted in 4 mL coupling buffer supplemented with 20 mM TCEP and incubated at room temperature with rotation for 30 minutes. The diluted Cys-OtUBD was loaded onto the SulfoLink resin, and the mixture was incubated at room temperature for 30-60 minutes with rotation. (Cys-His_6_-OtUBD required a longer incubation time (60 min) than Cys-OtUBD (30 min)) The resin was allowed to settle for another 30 minutes before being drained and washed once with 6 mL coupling buffer. 4 mL freshly prepared 50 mM L-cysteine dissolved in coupling buffer (pH adjusted to 8.5 with NaOH) was added to the resin and the mixture was incubated at room temperature for 30 minutes with rotation. The resin was allowed to settle for another 30 minutes before being drained and washed with 12 mL 1 M NaCl followed by 4 mL OtUBD column buffer (50 mM Tris•HCl, 150 mM NaCl, 1 mM EDTA, 0.5% Triton-X, 10% glycerol, pH 7.5). For long-term storage (more than 2 days), the resin was stored in column buffer containing 0.05% NaN_3_ and kept at 4°C. The resin could be stored at 4°C for up to 2 months without losing its efficiency. Longer storage times have not been tested.

#### Negative control Cys-coupled resin

The negative control resin was made by capping the reactive groups of the SulfoLink™ resin with cysteine following the manufacturer’s protocol. Specifically, 2 mL (bed volume) of resin was incubated with 4 mL freshly prepared 50 mM L-cysteine dissolved in coupling buffer (pH 8.5) at room temperature for 30 minutes with rotation. The resin was subsequently treated and stored as described above for the Cys-OtUBD resin.

#### TUBE resin

TUBE resin was made by conjugating Cys-His_6_-4xTR-TUBE to the SulfoLink resin following similar procedures as the OtUBD resin with some modifications. In particular, 2.33 mg of Cys-His_6_-4xTR-TUBE was diluted in 4 mL SulfoLink coupling buffer supplemented with 0.5 M guanidine•HCl and 20 mM TCEP. Guanidine was added to minimize precipitation of 4xTR-TUBE protein during incubation. Four mL of the diluted TUBE solution was added to 2 mL of SulfoLink™ resin and the mixture was incubated at room temperature for 1 hour with rotation. The rest of the preparation steps were the same as for OtUBD resin.

#### FK2 resin

FK2 anti-ubiquitin antibodies were covalently linked to a Protein-G resin following a previous protocol with modifications (20). Briefly, 500 μg of FK2 mouse monoclonal IgG1 antibody (Cayman Chemical) was diluted in 500 μL DPBS. The solution was added to 250 μL (bed volume) Protein G Sepharose™ 4 Fast Flow resin (GE Healthcare) prewashed with DPBS. The mixture was incubated at 4°C for 2 hours with rotation. The resin was washed twice with 100 mM triethanolamine•HCl (pH 8.3), and the antibody was then crossed-linked to the resin by incubation with 500 μL 50 mM dimethyl pimelimidate (DMP) dissolved in 100 mM triethanolamine•HCl buffer (pH 8.3) for 4 hours at 4°C with rotation. The reaction was terminated by incubating with 1.5 mL 100 mM Tris•HCl buffer (pH 7.5) for 2 hours at room temperature. Unconjugated antibody was removed from the resin by washing with 500 μL of 100 mM glycine•HCl buffer (pH 2.5). The resin was equilibrated with DPBS and stored at 4 °C before use.

## Ubiquitin-conjugate purifications with protein-linked resins

### Native conditions

#### Preparation of yeast lysate

For extraction of frozen yeast powder, 1 volume of cold native lysis buffer (50 mM Tris•HCl, 300 mM NaCl, 1 mM EDTA, 0.5% Triton-X100, 20 mM NEM, cOmplete mini EDTA-free protease inhibitor cocktail (Roche), 1 mM PSMF, pH 7.5) was added to extract proteins. The mixture was vortexed thoroughly and incubated on ice for 10 minutes with intermittent vortexing. The crude extract was centrifuged at 21,000 x *g* for 12 minutes, and the supernatant was carefully transferred to a clean tube.

Alternatively, yeast could be lysed by glass bead beating. Cell pellets were resuspended in 1 mL cold native lysis buffer containing 10% glycerol and 0.6 mL acid-washed glass beads (Sigma) and lysed in a FastPrep™ homogenizer (MP Bio) at 4°C (5.0 m/s, 3x(30 sec, 1 min rest on ice), 4 min rest on ice, 3x(30 sec, 1 min rest on ice)). The resulting mixture was left on ice for 5 more minutes and then centrifuged at 8000 x *g* for 5 minutes at 4°C. The supernatant was transferred to a new tube while 0.5 mL more lysis buffer was added to the beads and pelleted cell debris. The pellet was resuspended and treated as above. The supernatants were combined and centrifuged at 21,000 x *g* for 12 minutes at 4°C. The cleared lysate was transferred to a clean tube.

For mammalian cells, the frozen cell pellets were resuspended in cold native lysis buffer and incubated on ice for 30-40 minutes with occasional vortexing. After centrifugation at 21,000 x *g* for 20 minutes, clarified lysates were transferred to a fresh tube.

Protein concentration in the lysates was measured by the BCA assay, and lysates were adjusted to 2–4 mg/mL final protein concentration using native lysis buffer.

#### Pulldowns

A suitable amount of resin was either transferred to a disposable gravity column or, for smaller scale experiments, a microcentrifuge tube. Typically, 25 μL of resin (bed volume) was used for each 1 mg of lysate protein. For the proteomics experiments in this study, 0.25–1.6 mL of resin was used for each pulldown sample. (Here we describe the procedures used for gravity column-based experiments. For adaption to a microcentrifuge-based experiments, the users could pellet the resin at 1,000 x *g* for 1 min before removal of supernatant.) The storage buffer was drained, and the resin was equilibrated with 5 resin volumes of OtUBD column buffer. If the pulldown was performed for LC-MS/MS analysis, the resin was washed with 2 bed volumes of elution buffer (100 mM glycine•HCl, pH 2.5) and then immediately equilibrated by passing 20 bed volumes of column buffer through the resin.

Lysate was added to the equilibrated resin, the column was capped, and the mixture was incubated at 4°C for 2.5 hours with rotation. The resin was allowed to settle for 10 minutes, and the unbound solution was drained and collected as the flow-through. The resin was washed by passing 15 column bed volumes of column buffer, 15 volumes of Wash Buffer-1 (50 mM Tris•HCl, 150 mM NaCl, 0.05% Tween 20, pH 7.5) and 15 volumes of Wash Buffer-2 (50 mM Tris•HCl, 1 M NaCl, pH 7.5) sequentially through the resin.

#### Elution

If the downstream application was only Western blotting, the bound proteins could be eluted by incubating the resin with 2 to 3 resin volumes of 1x SDS sample buffer (50 mM Tris•HCl, pH 6.8, 2% SDS, 5% glycerol, 100 mM DTT, 0.005% bromophenol blue) for 15 minutes at room temperature with rotation. In our hands, the TUBE resin could only be efficiently eluted using this method.

If the purified proteins were to be analyzed by LC-MS/MS, 2 resin volumes of pure water were passed through the resin to push off residue buffers. Then, bound proteins were eluted by incubation in 2 resin volumes of elution buffer (100 mM glycine•HCl, pH 2.5) for 5 minutes at 4°C with rotation. The eluate was collected and immediately neutralized with 0.2 resin volume of 1M Tris•HCl pH 9 buffer. The elution process was repeated to ensure complete elution (the two eluates, E1 and E2, were sometimes combined to give eluate E). In some experiments, the first elution step was done with 100 mM glycine•HCl, pH 3.0 and a third elution step, also with the pH 2.5 buffer, was included to ensure complete elution.

Ubiquitin conjugates in the input, flow-through and eluate for each sample were analyzed by anti-ubiquitin Western blotting. The volume loaded onto the SDS-PAGE gel for each sample was normalized to reflect a 1:1:1 scaling of input, flow-through and eluate (e.g. if the total volume of the eluate is 1/10 that of the input, we load 1 volume of the input and 0.1 volume of the eluate on the same SDS-PAGE gel.) unless otherwise specified. Total protein from the pulldowns was analyzed by SYPRO Ruby staining of the gels. SYPRO™ Ruby stained gels were imaged on a Bio-Rad ChemiDoc imager and quantified using ImageJ software.

### Denaturing conditions

#### Preparation of lysates

Yeast or human cells were lysed as described above for the native condition protocol. After the measurement of protein concentration, the lysates were adjusted to up to 12.58 mg/mL protein with native lysis buffer. The lysate was kept on ice for the whole duration until appropriate amounts of solid urea were added directly to the native lysate to reach a final concentration of 8 M (1 g of urea was added per 0.763 mL of lysate; calculations were based on (48)), and the lysate was vortexed and agitated until the urea had fully dissolved. The urea lysate was incubated at 25°C for 30 minutes, chilled on ice and diluted 1:1 with native lysis buffer (final concentration of urea, 4 M).

Alternatively, in the experiment described in Fig. 4B (D2 condition), cells were lysed directly in urea lysis buffer (50 mM Tris•HCl, 300 mM NaCl, 8 M urea, 1 mM EDTA, 0.5% Triton-X, 20 mM NEM, cOmplete mini EDTA-free protease inhibitor cocktail (Roche), 1 mM PSMF, pH 7.5) by bead-beating. The concentration of the cleared lysate was determined by BCA assay and the concentration was adjusted to match other samples. The cleared lysate was incubated at 25°C for 15 minutes, chilled on ice and diluted 1:1 with native lysis buffer. This method could in theory include insoluble ubiquitylated proteins and may be useful in specific applications.

#### Pulldown and elution protocols

Pulldown procedures were similar to those described above under the native pulldown protocol except that the first wash step was done with column buffer containing 4 M urea. Elution steps are the same as described in the native pulldown protocol.

### M48 DUB treatment of yeast cell lysates

Yeast powder resulting from grinding the BY4741 strain in liquid nitrogen was reconstituted in M48 lysis buffer (50 mM Tris•HCl, 300 mM NaCl, 1 mM EDTA, 0.5% TritonX, 10% glycerol, pH 7.5, supplemented with 7.6 μM pepstatin A, 5 mM aminocaproic acid (ACA), 5 mM benzamidine, 260 μM AEBSF, 1 mM PMSF and 1 mM DTT), incubated for 10 minutes on ice and clarified by centrifugation at 21,000 x *g* at 4 °C. Inhibitors of cysteine proteases were avoided to prevent inhibition of the M48 cysteine protease (78). Protein concentrations in the lysates were determined by BCA assay, and the lysates were adjusted to 2 - 4 mg/mL protein with M48 lysis buffer. M48 DUB was added to the lysate to give a final enzyme concentration of 100 nM. The mixture was incubated at 37°C with rotation for 1 hour before subjecting to pulldown analysis. In the control samples where M48 was not added, 10 mM NEM and 20 μM MG132 (a proteasome inhibitor) were also included in the lysis buffer.

### Immobilized metal affinity chromatography (IMAC) under denaturing conditions

Eluates from the OtUBD pulldowns were denatured by adding a solid denaturant, either urea to a final concentration of 8 M or guadinine•HCl to a final concentration of 6 M. (Amounts of denaturants were calculated based on (48).) After the denaturant had fully dissolved, the solution was incubated at 25°C for 30 minutes before applying to a pre-washed HisPur™ Cobalt resin (Thermo Scientific). The mixture was incubated at room temperature with rotation for 1.5 hours, washed with 8 M urea wash buffer (50 mM Tris•HCl, pH 7.5, 8M urea), and eluted twice, each time by boiling for 5 minutes in 2 resin volumes of 500 mM imidazole in 2x SDS sample buffer (100 mM Tris•HCl, pH 6.8, 4% SDS, 10% glycerol, 200 mM DTT, 0.01% bromophenol blue). Samples were resolved by SDS-PAGE and analyzed by anti-ubiquitin immunoblotting and SYPRO Ruby staining. Specifically, samples containing guanidine was first diluted with 3 portions of pure H_2_O and then carefully mixed with 4x SDS sample buffer before loaded onto an SDS-PAGE gel to avoid precipitation of SDS.

### Proteomics

#### Sample preparation

Frozen samples were dehydrated on a lyophilizer (Labconco). For all samples except for those in the OtUBD/FK2 comparison experiment, the dry content was reconstituted in pure water and subjected to a methanol-chloroform extraction as described earlier (79).

#### In solution Protein Digestion

Protein pellets were dissolved and denatured in 8M urea, 0.4M ammonium bicarbonate, pH 8. The proteins were reduced by the addition of 1/10 volume of 45mM dithiothreitol (Pierce Thermo Scientific #20290) and incubation at 37°C for 30 minutes, then alkylated with the addition of 1/20 volume of 200mM methyl methanethiosulfonate (MMTS, Pierce Thermo Scientific #23011) with incubation in the dark at room temperature for 30 minutes. Using MMTS avoids the potential false positive identification of GG modification arising from iodoacetamide (IAA) alkylation (80). The urea concentration was adjusted to 2M by the addition of water prior to enzymatic digestion at 37°C with trypsin (Promega Seq. Grade Mod. Trypsin, # V5113) for 16 hours. Protease:protein ratios were estimated at 1:50. Samples were acidified by the addition of 1/40 volume of 20% trifluoroacetic acid, then desalted using BioPureSPN PROTO 300 C18 columns (The Nest Group, # HMM S18V or # HUM S18V) following the manufacturer’s directions with peptides eluted with 0.1% TFA, 80% acetonitrile. Eluted peptides were speedvaced dry and dissolved in MS loading buffer (2% acetonitrile, 0.2% trifluoroacetic acid). A nanodrop measurement (Thermo Scientific Nanodrop 2000 UV-Vis Spectrophotometer) determined protein concentrations (A260/A280). Each sample was then further diluted with MS loading buffer to 0.08µg/µl, with 0.4ug (5µl) injected for most LC-MS/MS analysis, except for the negative control samples, which were diluted to and injected the same volume as the corresponding OtUBD pulldown samples.

#### LC-MS/MS on the Thermo Scientific Q Exactive Plus

LC-MS/MS analysis was performed on a Thermo Scientific Q Exactive Plus equipped with a Waters nanoAcquity UPLC system utilizing a binary solvent system (A: 100% water, 0.1% formic acid; B: 100% acetonitrile, 0.1% formic acid). Trapping was performed at 5µl/min, 99.5% Buffer A for 3 min using an ACQUITY UPLC M-Class Symmetry C18 Trap Column (100Å, 5 µm, 180 µm x 20 mm, 2G, V/M; Waters, #186007496). Peptides were separated at 37°C using an ACQUITY UPLC M-Class Peptide BEH C18 Column (130Å, 1.7 µm, 75 µm X 250 mm; Waters, #186007484) and eluted at 300 nl/min with the following gradient: 3% buffer B at initial conditions; 5% B at 2 minutes; 25% B at 140 minutes; 40% B at 165 minutes; 90% B at 170 minutes; 90% B at 180 min; return to initial conditions at 182 minutes. MS was acquired in profile mode over the 300-1,700 m/z range using 1 microscan, 70,000 resolution, AGC target of 3E6, and a maximum injection time of 45 ms. Data dependent MS/MS were acquired in centroid mode on the top 20 precursors per MS scan using 1 microscan, 17,500 resolution, AGC target of 1E5, maximum injection time of 100 ms, and an isolation window of 1.7 m/z. Precursors were fragmented by HCD activation with a collision energy of 28%. MS/MS were collected on species with an intensity threshold of 1E4, charge states 2-6, and peptide match preferred. Dynamic exclusion was set to 30 seconds.

#### Peptide Identification

Data was analyzed using Proteome Discoverer software v2.2 (Thermo Scientific). Data searching is performed using the Mascot algorithm (version 2.6.1) (Matrix Science) against a custom database containing protein sequences for OtUBD as well as the SwissProt database with taxonomy restricted to *Saccharomyces cerevisiae* (7,907 sequences) or *Homo sapiens* (20387 sequences). The search parameters included tryptic digestion with up to 2 missed cleavages, 10 ppm precursor mass tolerance and 0.02 Da fragment mass tolerance, and variable (dynamic) modifications of methionine oxidation; N-ethylmaleimide, N-ethylmaleimide+water, carbamidomethyl, or methylthio on cysteine; and GG adduct on lysine, protein N-terminus, serine, threonine or cysteine. Normal and decoy database searches were run, with the confidence level set to 95% (p<0.05). Scaffold (version Scaffold_5.0, Proteome Software Inc., Portland, OR) was used to validate MS/MS based peptide and protein identifications. Peptide identifications were accepted if they could be established at greater than 95.0% probability by the Scaffold Local FDR algorithm. Protein identifications were accepted if they could be established at greater than 99.0% probability and contained at least 2 identified peptides. GG modified peptides were further analyzed using Scaffold PTM 3.3 software.

Quantitative analysis was done by Scaffold 5 (Proteome Software) based on normalized total TIC (MS/MS total ion current). Pearson correlation coefficients were calculated using GraphPad Prism 9 software. Volcano plots were generated using GraphPad Prism 9 software. Proteins are selected as a potential E3 substrate if they meet one of the following criteria: 1) Its average quantitative value (normalized total TIC) is at least 1.5 times higher in the WT samples compared to the E3 deletion samples and p value < 0.05. 2) It appeared in at least 3 of the 6 technical replicates of the WT samples but not in any of the 6 technical replicates of the E3 deletion samples.

GO enrichment analysis on specific protein populations was performed using the online Gene Ontology engine (81–83) accessible at: http://geneontology.org/.

## Acknowledgements

We thank Dr. Chin Leng Cheng and Dr. Hong-Yeoul Ryu for providing yeast strains used in the manuscript. We thank the Schlieker lab at Yale University for sharing of the M48 DUB expression plasmid, tissue culture space, and equipment. We thank the Paulsen lab Yale University for sharing of their ChemiDoc instrument.

## Supplementary Figure Legends

**Figure S1**

A, B. Ubiquitin pulldowns with different amounts of MBP-OtUBD. In A, the pulldown was performed by first binding MBP-OtUBD to an amylose resin and then incubating the resin with yeast cell lysate. In B, pulldown was performed by incubating the lysate with MBP-OtUBD and then binding the complexes to amylose resin. U: unbound fraction; E: fraction eluted with maltose.

C. Anti-ubiquitin blot of MBP-OtUBD pulldowns from HEK293T whole cell lysates. U: unbound fraction; B: bound fraction (eluted with SDS sample buffer).

D. SYPRO Ruby protein stain of the eluates from OtUBD-resin in Fig. 2B. E1/E2/E3: eluted fractions from serial low pH elutions.

**Figure S2**

A. Western blots of yeast proteasomal subunits in OtUBD pulldown samples. Unmodified yeast proteasomal subunits (Rpt4, Rpt5, Pre6) bound to the OtUBD resin under native conditions but not following denaturation of the lysate prior to pulldown. N: Native condition; D: Denaturing condition.

B. Western blots of human proteasomal subunits in OtUBD pulldown samples. Unmodified human proteasomal subunit Rpt6 and Rpt4 bound strongly to OtUBD resin under native conditions but only weakly following extract denaturation. Modified Rpt6 and Rpt4 (likely ubiquitylated) bound to the OtUBD resin under both native and denaturing conditions. N: Native condition; D: Denaturing condition.

**Figure S3**

A. Yeast lysates analyzed by anti-ubiquitin blotting following their purification on TUBE and OtUBD resins. IN: input; FT: flowthrough; E: eluted fraction (SDS sample buffer elution); T: TUBE pulldown; U: OtUBD pulldown.

B. OtUBD pulldowns of ubiquitylated RNAPII subunit Rpb1 from human (HeLa) whole cell lysates under native conditions.

**Figure S4**

A. Representative anti-ubiquitin Western blot of OtUBD pulldowns (under native conditions) used for proteomics analysis. IN: input; FT: flowthrough; E1/E2/E3: eluted fractions from a series of low pH elutions.

B. Representative anti-ubiquitin blot of OtUBD pulldown samples following extract denaturation (urea) and used for proteomics analysis.

C. Representative SYPRO Ruby protein stain of OtUBD eluates resolved by SDS-PAGE.

D. Number of proteins detected in each biological replicate of OtUBD pulldown-MS and negative control. Error bar represents difference among technical replicates.

E. Box and whisker plot showing distribution of the quantitative value - total TIC (total ion current) values for each protein in each biological sample. Proteins in the negative controls generally present at much lower level compared to the OtUBD pulldown samples.

F. Adjusted number of proteins detected in each biological replicate of OtUBD pulldowns. Only proteins whose TIC value are at least 20 times higher in the OtUBD pulldown samples compared to the corresponding negative control samples are included.

G. Anti-ubiquitin Western blot of OtUBD pulldowns, and FK2 antibody IPs used for proteomics analysis. IN: input; FT: flowthrough; E: pooled eluted fractions; E1/E2: eluted fractions from a series of low pH elutions.

H. Quantitation (estimated based on total spectral counts) of different ubiquitin linkages under native and denaturing (urea) conditions in the BY4741 yeast ubiquitylome.

**Figure S5**

A. Representative anti-ubiquitin blot of OtUBD pulldowns from WT, *bre1△* and *pib1△* yeast lysates used for proteomics analysis. IN: input; FT: flowthrough; E: pooled eluted fractions.

B. Representative SYPRO Ruby gel showing the total proteins in eluates from OtUBD pulldown from WT, *bre1△* and *pib1△* yeast lysates.

C, D. Pearson correlation coefficients were calculated between each sample in the analyzed groups using normalized total TIC. Because ubiquitin is present in exceptionally high levels compared to all other proteins, it was excluded from the dataset for this analysis. With the exception of one *pib1Δ* sample, correlations between different samples were generally high, as expected if the majority of the ubiquitylome was not affected by deletion of a single E3. The low correlation in the single *pib1Δ* sample was likely due to an error during sample preparation, so the results were excluded from the quantitation.

**Figure S6**

A. Representative MS/MS spectrum of the Htb2 K111GG peptide.

B. Representative MS/MS spectrum of the Htb2 K123GG peptide. In addition to a/b/y ions, we identified multiple peaks from internal fragmentation and dehydration, potentially due to the serine/threonine-rich nature of the sequence.

C. Representative MS/MS spectrum of the YMR160W T534GG peptide.

## Supplementary Documents

Supplementary Data 1: List of plasmids and yeast strains used in this study.

Supplementary Data 2: Proteomics data in this study.

